# Quantitative cross-linking/mass spectrometry using isotope-labeled cross-linkers and MaxQuant

**DOI:** 10.1101/055970

**Authors:** Zhuo A. Chen, Lutz Fischer, Jüergen Cox, Juri Rappsilber

## Abstract

The conceptually simple step from cross-linking/mass spectrometry (CLMS) to quantitative cross-linking/mass spectrometry (QCLMS) is compounded by technical challenges. Currently, quantitative proteomics software is tightly integrated with the protein identification workflow. This prevents automatically quantifying other m/z features in a targeted manner including those associated with cross-linked peptides. Here we present a new release of MaxQuant that permits starting the quantification process from an m/z feature list. Comparing the automated quantification to a carefully manually curated test set of cross-linked peptides obtained by cross-linking C3 and C3b with BS^3^ and isotope-labeled BS^3^-d4 revealed a number of observations: 1) Fully automated process using MaxQuant can quantify cross-links in our reference dataset with 68% recall rate and 88% accuracy. 2) Hidden quantification errors can be converted into exposed failures by label-swap replica, which makes label-swap replica an essential part of QCLMS. 3) Cross-links that failed during automated quantification can be recovered by semi-automated re-quantification. The integrated workflow of MaxQuant and semi-automated assessment provides the maximum of quantified cross-links. In contrast, work on larger data sets or by less experienced users will benefit from full automation in MaxQuant.

**Abbreviations:**

BS^3^: Bis[sulfosuccinimidyl] suberate
CLMS: Cross-linking/mass spectrometry
MS1: the initial mass-to-charge-ratio (m/z) spectrum collected for all components in a sample.
QCLMS: Quantitative cross-linking/mass spectrometry

## Introduction

The function of proteins is often linked to conformational rearrangements. Quantitative cross-linking/mass spectrometry (QCLMS) using isotope-labeled cross-linkers (1–4) is emerging as a new strategy to study such conformation changes of proteins (5). Applications include the trans-membrane protein complex F-type ATPases (6), the multidomain protein C3 converting into C3b (7), modelling the structure of iC3 (Chen *et al.*, MCP/2015/ 056473) and the maturation of the proteasome lid complex (8). These show that the QCLMS approach has great potential for detecting protein conformational changes in macro protein assemblies and possibly also complex protein mixtures such as large protein networks. However, great challenges result from the size and complexity of datasets generated when studying such large and complex protein systems.

Manually interrogating QCLMS data (6, 9) by experts can be superior to the performance of automated algorithms, however it is also time consuming, subject to human handling errors and invites the omission of important controls. Consequently, a benchmark study (7) relied on a semi-automated quantitation setup for cross-linking data by exploring the functionality of a quantitative proteomics software Pinpoint (Thermo Scientific). However, still, manually inspecting and correcting quantitation results from Pinpoint was tedious, required expertise and will become increasingly impractical as data size increases. Recently, Kukacka et al. presented a workflow using mMass at the example of calmodulin (17 kDa) in presence and absence of Ca^2+^ (10). However, the scalability of this approach remains to be shown. As a prove-of-principle, we established a computational workflow to quantify the signals of cross-linked peptides in an automated manner (5). We developed an elementary computational tool, XiQ (5), which allowed us to accurately quantify our model dataset. Yet, XiQ has three major drawbacks: 1) it is not optimized for chromatographic feature detection; 2) XiQ is a command line based application and lacks an easy user interface; 3) XiQ does not visualize its output and hence does not facilitate manual inspection and validation.

To overcome these disadvantages, we exploited the well-established chromatographic feature detection function and user friendly interface of one of the most commonly used quantitative proteomics software tools, MaxQuant (11). While developed originally for the analysis of SILAC data (12) MaxQuant has undergone recent expansion of workflows, including label-free quantitation (13) and widening its vendor support (14). Based on our initial assessment of MaxQuant's weaknesses in the context of QCLMS (5), we developed here a new version of MaxQuant for carrying out automated quantitation in cross-link experiments (Fig.1). We generated a reference dataset, based on our benchmark QCLMS analysis of C3 and C3b (7), to test the performance of this and future new tools. The results showed that experiments with replicated analysis and label-swap provided effective quality control for fully automated quantitation. Finally, we suggest an integrated workflow of MaxQuant and semi-automated processing. Pinpoint provides a platform for validating and correcting fully automated quantitation results, improving both data recall rate and quantitation accuracy.

**Figure 1:**
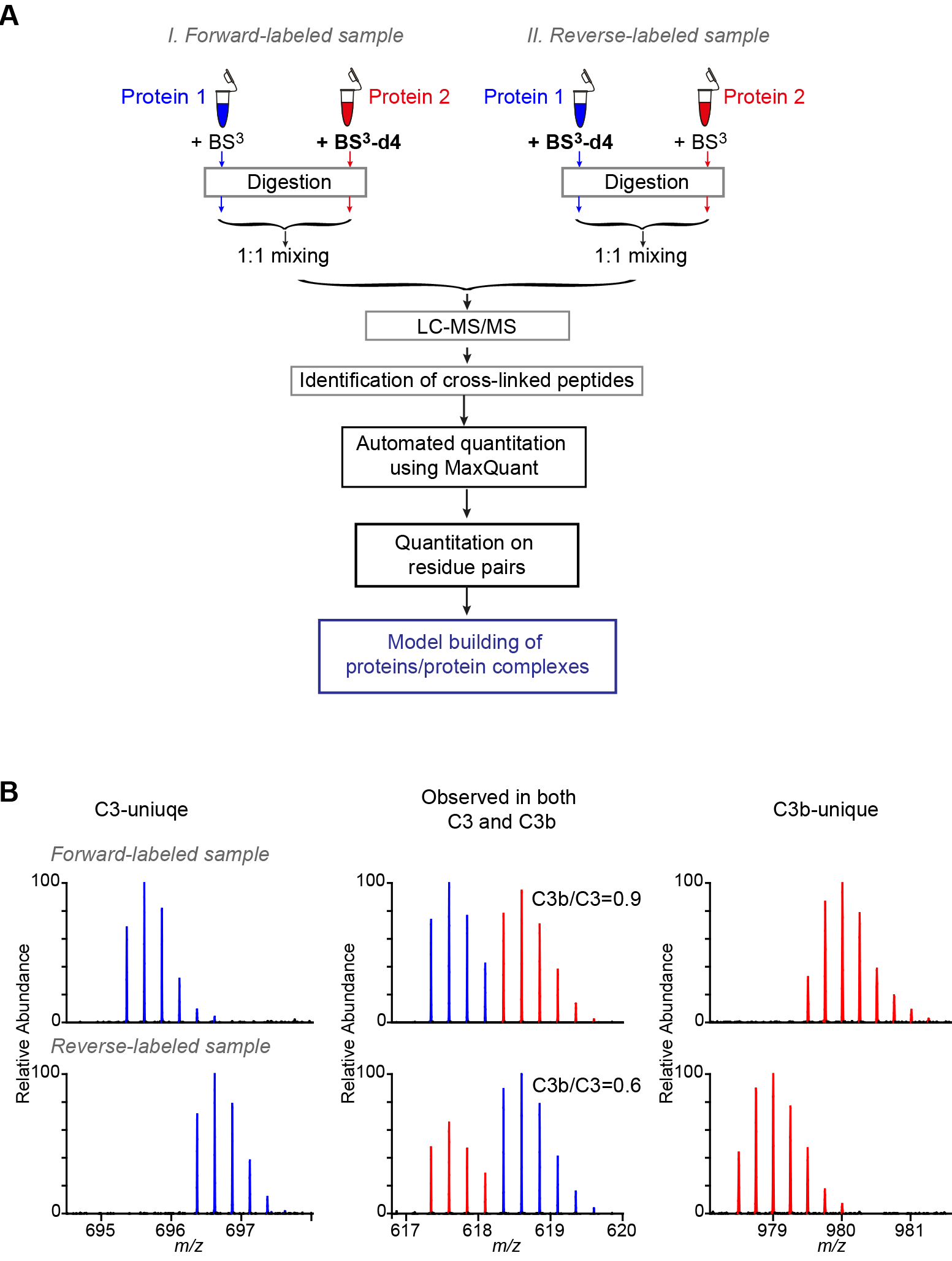
Automated quantitation for cross-linked peptides using MaxQuant. (**A**) The workflow of automated quantitation for cross-linked peptides using MaxQuant.(**B**) Example mass spectrometric signals of C3-unique (left), C3b-unique (right) and C3-C3b common (middle) cross-linked peptides in both forward-labeled and reverse-labeled analysis.

## Experimental Procedures

### Datasets

Dataset 1 was published previously (5). It comprised nine LC-MS files containing data on isotopically mixed, cross-linked human serum albumin (HSA). HSA was cross-linked with mixtures of bis[sulfosuccinimidyl] suberate-d0 (BS^3^) (Thermo Fisher Scientific) and its deuterated form bis[sulfosuccinimidyl] 2,2,7,7–suberate-d4 (BS^3^–d4) (Thermo Fisher Scientific). For the purpose of quantitation, BS^3^ and BS^3^-d4 were mixed with three molar ratios, 1:1, 1:2 and 1:4 (with three replicas for each ratio). Cross-linked HSA was then digested by trypsin and analyzed by LC-MS/MS using an LTQ Orbitrap Velos instrument (Thermo Scientific) as described (5). This dataset revealed weaknesses of a previous version of MaxQuant (version 1.2.2.5) in quantifying cross-linked peptides (5). The dataset was used again, to assess if the previously observed problems where successfully addressed using the here described new version of MaxQuant (version 1.5.4.1).

Dataset 2 was established here based on our benchmark QCLMS analysis on complement protein C3 *versus* its active product C3b (7). It therefore constitutes a more real analysis situation of an actual conformation change. Mass spectrometric raw data is available in the ProteomeXchange Consortium (15)

(http://proteomecentral.proteomexchange.org) via the PRIDE partner repository with the dataset identifier PXD001675. Peak lists were generated using MaxQuant version 1.2.2.5 (11) with default parameters, except that “Top MS/MS Peaks per 100 Da” was set to 20. The peak lists were searched against C3 and reversed C3 sequences (as decoy) using Xi software (ERI, Edinburgh) for identification of cross-linked peptides.Search parameters were as follows: MS accuracy, 6 ppm; MS2 accuracy, 20 ppm; enzyme, trypsin; specificity, fully tryptic; allowed number of missed cleavages, four; cross-linker, BS^3^/BS^3^-d4; fixed modifications, carbamidomethylation on cysteine; variable modifications, oxidation on methionine, modifications by BS^3^/BS^3^-d4 that are hydrolyzed or amidated on the other end. The reaction specificity of BS^3^ for modification and crosslinking was assumed to be lysine, serine, threonine, tyrosine and protein N-termini. The identified cross-linked peptides were quantified based on their precursor MS signals. Quantitation was carried out with a semi-automated workflow using Pinpoint software (Thermo Fisher Scientific) (7)

103 cross-linked peptides (Supplemental Table S1) that were included in the model dataset fulfilled two major criteria: 1) each cross-linked peptide was reproducibly and consistently identified and quantified in both label-swap replicas. 2) Conformational dynamic information carried by the quantitation results of these cross-linked peptides had been orthogonally validated by crystal structures of C3 and C3b. Following key identification information of these 103 cross-linked peptides in both forward-labeled and reverse-labeled analysis were used for constructing input for MaxQuant based quantitation: m/z, charge state, retention time, labeling status (BS^3^ cross-linked or BS^3^-d4 cross-linked), mass of the isotope label (4.02511 Da for BS^3^-d4), number of isotope labels.

### Quantitation of cross-link data using Maxquant software

Quantitation of cross-link data using a new release of MaxQuant (version 1.5.4.1 http://www.coxdocs.org/doku.php?id=maxquant:common:download_and_installation) was evaluated using the reference datasets described above. A library file (feature list) was constructed using Microsoft Excel, based on identification results of cross-linked peptides, for each model dataset and served as input file for peptide identities. The feature list for dataset 2 was shown in as an example in Supplemental Table S2 as an example. To carry out quantitation (Fig 2), all involving raw mass spectrometric data files were loaded. Under “Group-specific parameters”, for the “General” parameters, “Quantification only” mode was selected from the “type” options; for the “Advanced” parameters, “Match from file” was picked and the library file was then loaded. “Mass tolerance” was set to 6 ppm, “Time tolerance” was set to 3 minutes and “Time tolerance for label” was set to 2 minutes. The automated quantitation results were written into the “libraryMatch.txt” file in the folder “combined”.

**Figure 2:**
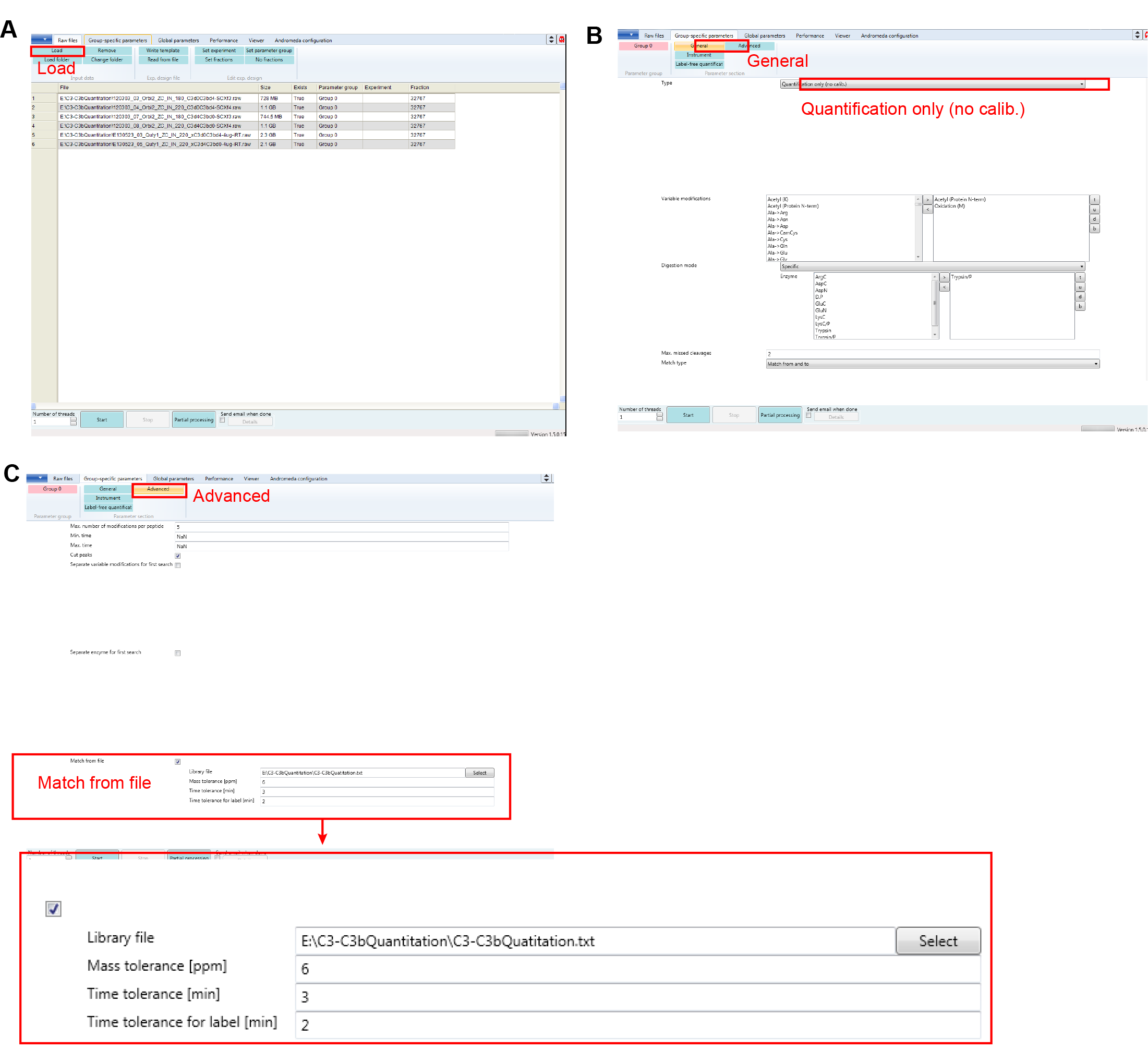
Quantitation using a feature list in a new release of MaxQuant. Three key steps for setting up a MaxQuant quantitation for cross-linked peptides and other analytes currently not native to the MaxQuant identification workflow in the new user interface of MaxQuant (version 1.5.4.1): (**A**) Loading raw data. (**B**) Selecting identification independent quantitation. (**C**) Uploading peptide library and define chromatography related parameters.

The output of MaxQuant quantitation was subsequently processed using Microsoft Excel. Quantitation was summarized first within each quantitation samples (forward-labeled or reverse-labeled). Quantitation results of a cross-linked peptide at different charge states were summarized as intensity weighted average. Quantitation of each cross-linked residue paired is summarized as the median of quantitation results of all its supporting cross-linked peptides. A cross-linked residue pair is determined as a C3/C3b-uique cross-link only if all its supporting cross-linked peptides are quantified as C3/C3b-unique signals accordingly. Quantitation of both cross-linked peptides and cross-linked residue pairs in the label-swap replicas were also compared on both signal type assignments and C3b/C3 signal ratios for further confirmation of the quantitation conclusion.

### Accession codes for review

The mass spectrometry proteomics data for dataset 1 and dataset 2 have been deposited to the ProteomeXchange Consortium (15)(http://proteomecentral.proteomexchange.org) via the PRIDE partner repository.

#### Dataset 1

Dataset identifier PXD004107

Project Webpage:http://www.ebi.ac.uk/pride/archive/projects/PXD004107FTP Download:ftp://ftp.pride.ebi.ac.uk/pride/data/archive/2016/05/PXD004107

#### Dataset 2

Dataset identifier PXD001675.

## Results and Discussions

### Automated quantitation for cross-linked peptides using Maxquant

As one of the most commonly used quantitation software tools for proteomics studies, MaxQuant has a well-established algorithm for chromatographic feature detection. It also provides a user-friendly interface. Recently, we tested the possibility of quantifying crosslink data by adapting the standard MaxQuant workflow. To obtain a model dataset, named here dataset 1, HSA was cross-linked with a mixture of BS^3^ and BS^3^-d4 in difference mixing ratios. This generated doublet MS signals for each cross-links with light to heavy signal ratios of 1:1, 1:2 and 1:4, respectively (5). Unfortunately, we found that the routine quantitation algorithm for SILAC-based studies in MaxQuant was not suitable for QCLMS analysis (5). The isotope effect of deuterium in the labeled cross-linker often led to shifts in retention time for the normal (light) compared to the heavy version of a cross-linked peptide. Such retention shift hindered MaxQuant from providing accurate quantitation for cross-links (Fig. 3A and B) (5). In addition, MaxQuant did not allow us to specify a feature list for quantitation and thus to direct the software towards the MS1 features of interest to us.

**Figure 3:**
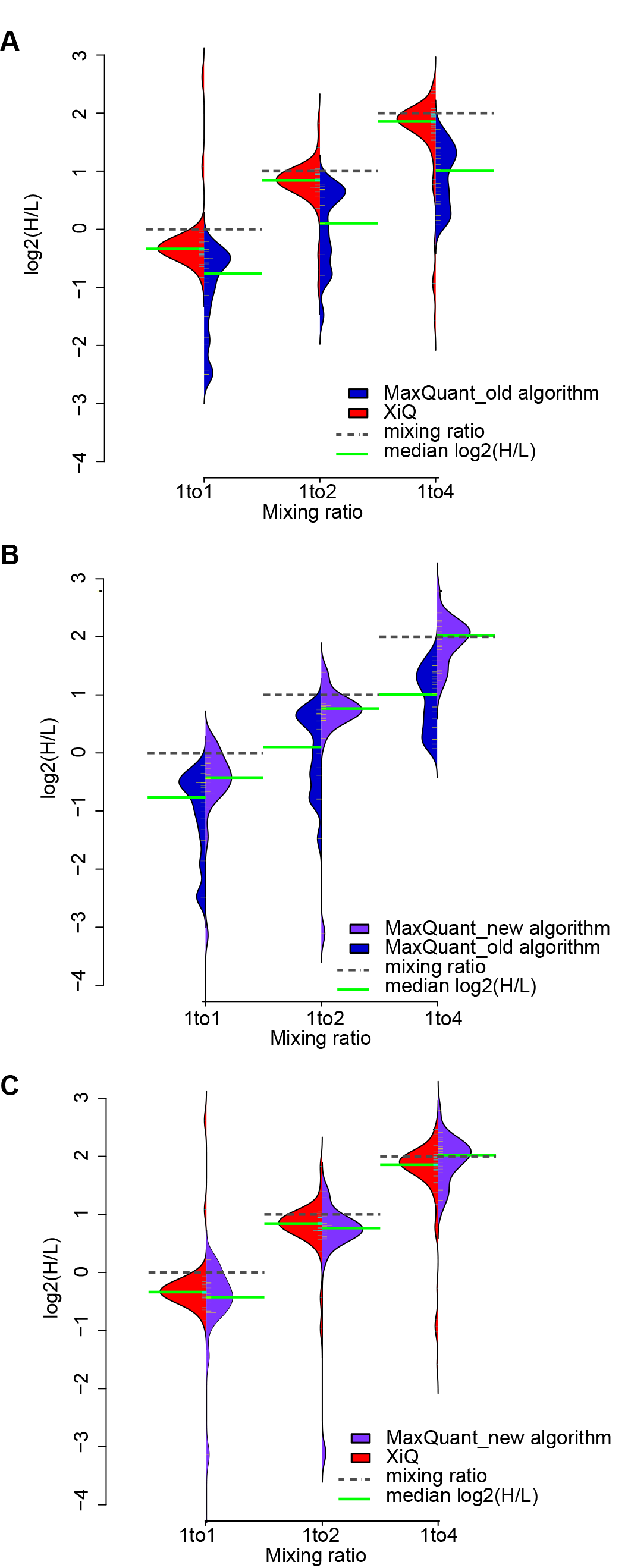
MaxQuant is capable of quantifying cross-linked peptides. Quantitation results of dataset 1 using quantification from feature list in MaxQuant are compared with the previously published results using standard quantification in MaxQuant and XiQ. (**A**) Bean plots showing the distribution of H/L ratios of cross-links for each mixing ratio quantified using XiQ (red) and standard quantification in MaxQuant (blue). (**B**) Bean plots showing the distribution of H/L ratios of cross-links for each mixing ratio quantified using quantification from feature list in MaxQuant (purple) and standard quantification in MaxQuant (blue). (**C**) Bean plots showing the distribution of H/L ratios of cross-links for each mixing ratio quantified using quantification from feature list in MaxQuant (purple) and XiQ (red).

Here we developed a new version of MaxQuant (version 1.5.4.1) with two major new features to enable quantitation of cross-linking data. 1) A new “Quantification only” mode is available in the user interface that allows for quantitation independent of the identification module, therefore enables quantitation of cross-links and other signals currently not native to the MaxQuant identification workflow. 2) Furthermore, the quantitation algorithm used in such cases builds on the same quantitation workflow that we have established in XiQ, quantifying the peaks of a doublet separately and then forming their intensity ratio, instead of the traditional approach of calculating a ratio for each MS1 scan and then taking the median. As a first step to provide quantitative information, MaxQuant requires the m/z and elution time of MS1 features to be quantified. Normally, this information is forwarded internally from Andromeda (16), the peptide identification module of MaxQuant. However, Andromeda is currently incapable of identifying linked peptides and hence an alternative route has to be taken. Instead, cross-linked peptides are imported from a library file (feature list), provided by the user (Fig. 2, Supplemental Table 2). As a positive side aspect, the user can choose freely among the available software tools for generating peak-lists and searching databases to identify cross-linked peptides in order to construct the input library file in.txt format. The following columns are required in this txt file: “run_name”, “precursor_charge”, “ms2 retention time”, “precursor_mz”, “number of cross-linker” and “cross-linker”. Additional information can also be included in the file to facilitate subsequent data processes (Supplemental Table 2). For each entry in the feature list, MaxQuant first identifies its precursor chromatographic feature in the raw data and extracts the intensity of the elution peak. In addition, MaxQuant allocates the doublet partner based on the mass difference between the normal and heavy forms of the peptide and extracts the partner's signal intensity. In the output of MaxQuant, intensities of both light and heavy signals are listed and heavy/light signal ratios are calculated for each entry. The output file from MaxQuant is in text format and can be further processed using spreadsheet applications such as Microsoft Excel (Experimental procedures).

Independent read-out of intensities of paired light and heavy signals minimized the impact of retention time shifts on final signal ratios (5). Testing with dataset 1, described above, showed that the new implementation of MaxQuant has a significantly improved performance on quantifying cross-links and is now capable of quantifying cross-linking data. While MaxQuant using its standard algorithm quantified cross-links clearly less well than XiQ (Fig. 3A) this improved when implementing the XiQ workflow in MaxQuant (Fig. 3B, C). Now both programs show comparable quantification success for cross-linked peptides, albeit with many other benefits being unique to MaxQuant.

### A reference dataset for evaluating performance of quantitation tools for QCLMS analysis

To test the ability of the MaxQuant to conduct an actual QCLMS analysis, we generated a reference list of quantified cross-linked peptide pairs from our benchmark study of complement protein C3 and its active form C3b. This dataset is referred to here as dataset 2. In our benchmark study, the structures of C3 and C3b were compared using QCLMS and cross-linked peptides were quantified using a semi-automated approach based on Pinpoint software (Thermo Scientific) (7). PinPoint largely facilitated the manual quantification process. Without manual assistance, the software is not capable of providing reliable data as it frequently errors in the selection of elution peaks. In this way we ensured the highest possible quantification quality with the current technology. The available crystal structures allowed to confirm our success of cross-link identification and the sorting into unique for one protein (singlet) or shared between C3 and C3b (doublet) (7).

We used the entire available raw data of that study but selected a subset of quantified cross-linked peptide pairs. The data comprised four data sets, a pair of forward and reverse label experiments acquired on a Q Exactive mass spectrometer (Thermo Fisher) and a second pair of forward and reverse label experiments acquired on an LTQ Orbitrap Velos mass spectrometer (Thermo Fisher). MaxQuant does currently not allow match between runs for cross-link data. Therefore, we included only unique peptide pairs that were identified and quantified in one complete experiment, i.e. in matching forward and reverse label experiments acquired on the same mass spectrometer. We arrived at 103 unique, cross-linked peptide pairs (Supplemental Table S1). These contained 59 unique linked residue pairs, including 16 (34 peptide pairs) unique to C3, 14 (17 peptide pairs) unique to C3b and 29 (52 peptide pairs, C3b/C3 signal ratios range 0.1-71.8) shared by both (detected as doublet signals) (Fig. 1B). A total of 610 fragmentation spectra matched to our reference set of 103 unique, crosslinked peptide pairs.

Such a dataset with our manually curated reference list of quantified features provides a high-quality test set for evaluating the performance of our new MaxQuant release. The raw data we used alone or together with our identifications and quantifications (Supplemental Table S1, S2) also offer the possibility to use them as a reference dataset for testing any QCLMS data processing setup. The raw data are publicly available via the ProteomeXchange (PRIDE) repository, dataset identifier PXD001675.

### Label-swap replica expose quantitation error in fully automated quantitation

We applied automated MaxQuant quantitation in a workflow that is equivalent to what has been applied in our benchmark study (Fig. 1A) and compared MaxQuant results against our carefully curated reference list of quantified features. Of 103 cross-linked peptides in the reference list, 92 were quantified using the automated MaxQuant process.

The 92 cross-linked peptides quantified by MaxQuant could be divided into three different sub-groups (Fig. 4A). 65 cross-linked peptides (150 quantified features, note that we combined multiple charge states into a single feature) were quantified by MaxQuant consistently in our replica with label-swapping. 11 cross-linked peptides (11 quantified features) were only quantified in a single replica. Moreover, 16 cross-linked peptides (36 quantified features) showed conflicting assignment into singlet versus doublet across two replicas. For a more direct comparison, the MaxQuant results were compared against the reference quantitation results on each individual quantified feature. We looked at signal type assignment (singlet versus doublet) and C3b/C3 signal ratios.

**Figure 4:**
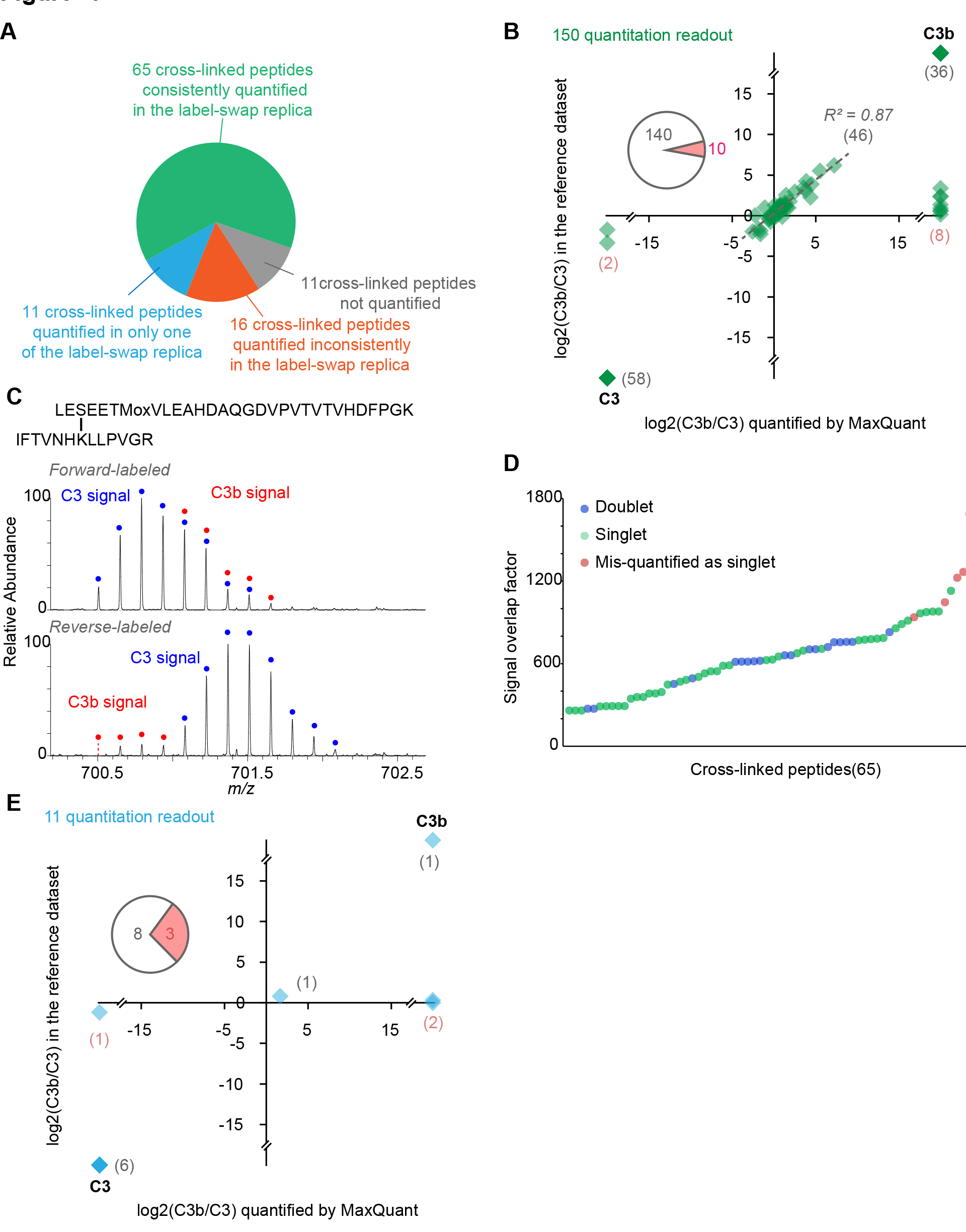
Label-swap replica expose quantitation error in fully automated quantitation. (**A**) 103 cross-linked peptides from the model dataset can be divided into four subgroups based on fully automated MaxQuant quantitation. (**B**) Cross-linked peptides that were consistently quantified in label-swap replica showed good agreement on both signal type assignment and C3b/C3 signal ratios between the MaxQuant quantitation and the reference dataset for 159 corresponding quantitation readout. (**C**) Implying from the MS signal of cross-linked peptide LES(cl)EETMoxVLEAHDAQGDVPVTVTVHDFPGK-IFTVNHK(cl)LLPVGR, it was mis-quantified as a C3 unique signal in the forward-labeled analysis (C3-light; C3b-heavy) due to highly overlapped isotope envelopes of the light and heavy signals, and in the reverse-labeled analysis (C3-heavy, C3b-light) due incomplete isotope envelope for the light signal. (**D**) All five cross-linked peptides that were mis-classified as singlet signals in both label-swap replicas showed larger overlap between their light and heavy signals in respect to correctly quantified doublet crosslinked peptides. The signal overlap factor is calculated as the mass of the cross-linked peptide divided by the mass difference between the light and heavy signals. (**E**) Comparison of MaxQuant quantitation readout against the reference dataset on signal type assignments and C3b/C3 signal ratios for 11 cross-linked peptides that were quantified in only one of the label-swap replicas.

The results of those 65 cross-linked peptides that were reproducibly quantified in replica by MaxQuant matched closely our reference data (Fig. 4B). MaxQuant succeeded in returning the expected classification (singlet or doublet) for 60 (92%) of these 65 cross-linked peptides and failed for 5 (8%). 140 out of 150 (93%) quantified features agreed with our reference data on signal type assignment. All 94 singlets were correctly classified. 46 doublet signals were also correctly classified and their C3b/C3 signal ratios show reproducibility of R^2^=0.87 between fully automated (MaxQuant) and previous manually curated quantitation. The remaining variation presumably resulted from differences in chromatographic feature detection between MaxQuant and Pinpoint. Surprisingly, MaxQuant misassigned 10 doublet signals (of 5 cross-linked peptides) as singlets. Doublet signals were repeatedly missed as a result of incomplete isotope envelope (e.g. missing mono-isotopic peak) which in turn was the result of low signal intensity and large peptide mass (Fig. 4C). A second reason was found in heavily overlapping isotope clusters of light and heavy signals. All five cross-linked peptides showed larger overlap between their light and heavy signals in respect to correctly quantified doublet cross-linked peptides. The signal overlap factor is calculated as the mass of the cross-linked peptide divided by the mass difference between the light and heavy signals (Fig. 4C, D).

For 11 cross-linked peptides, MaxQuant returned values only for one replica. This included seven singlets and one doublet (73%) in full agreement with our reference list. Three doublets (27%) were falsely called by MaxQuant as singlets (Fig. 4E). Because of this generally poorer quantitation accuracy (compared to 8% error when MaxQuant agreed in its call across replica) we did not include these peptides in the results of the automated analysis and re-quantified them manually in our expanded workflow. The same was done with those 16 cross-linked peptides where MaxQuant had conflicting signal type assignments from replica.

This detailed assessment led us to four conclusions. (1) Cross-linked peptide features are challenging to quantify. Reasons are largely weak signals due to sub-stoichiometric presence of cross-links, small mono-isotopic peak due to large peptide size, overlapping isotope clusters and retention time shift due to isotope effects. These problems are presumably inherent to cross-link studies and might be difficult to overcome experimentally. For example, when increasing the mass of the isotope-label to reduce peak-overlap one risks worsening the isotope effect on retention time. Further improvements in quantitation algorithms might play a larger role. (2) Nevertheless, MaxQuant reliably quantified a large subset of MS signals of cross-linked peptides. (3) Replicated analysis with label-swap provides an efficient quality control. Consistent quantitation in both label-swapping samples reveals accurate results and highlights problematic MS1 features. (4) Finally, cross-linked peptides that are not consistently quantified in both label-swapping replica by MaxQuant need to be reviewed individually or discarded.

Importantly, besides cross-linked peptides and MS1 features there is another level of information: residue pairs. Residue pairs are the information used subsequently in structural studies. Therefore, we combined data of cross-linked peptides into crosslinked residue pairs. For a residue pair to be assigned as unique to one conformation required that all its supporting cross-linked peptides were unique to this conformation. We argue that missing one partner of a low-intensity doublet is easily done. So, seeing even a single doublet for a residue pair suffices to shade sufficient doubt on an overall singlet assignment.

The 65 cross-linked peptides that were consistently quantified in label-swapping replica (see above) gave rise to 40 unique residue pairs and 35 (88%) of them were correctly recognized as singlet or doublet, respectively. The five misclassified residue pairs were each supported by a single cross-linked peptide, each observed by a single MS1 feature in the two replicas. In consequence, it appears prudent to base automated quantitative cross-linked residue pair data always on multiple quantified features. Arguably, also manual quantitation requires great care when basing arguments on a single feature. Requiring multiple features is longstanding practice when working with proteins in quantitative proteomics. Both, proteins and cross-linked residue pairs are quantified based on peptide signals. A fundamental difference between the two is, however, that usually much fewer observations are combined to give a value for a crosslinked residue pair (here in average 150/40 =3.8) than for a quantified protein (for example, in a chromatin study of our lab this was 11.7(17)). This limits quantification accuracy and recall rates for linked residue pairs.

### An integrated quantitation workflow for cross-linking data

Consistent quantitation in label-swap replicas ensures accurate quantitation. However, relying here solely on automated data processing reduces the recall rate. To make the most out of the available data requires manual assessment and correction of problematic m/z features. While MaxQuant provides a fast route for non-problematic features and also for spotting problematic ones it does not currently provide a platform for manually interrogating peaks. This led us to an integrate workflow for QCLMS analysis (Fig. 5A):

1. QCLMS analyses must be conducted in replicas and label-swapping.
2. Identified cross-linked peptides are quantified using MaxQuant in a fully automated manner based on their MS signals. The identification information of cross-linked peptides are imported to MaxQuant in format of a peptide library (feature list). This is an open and flexible entry point, which is independent of the algorithm used for identifying cross-linked peptides. In fact, MaxQuant can quantify peptides from any other source now in this way.
3. Cross-linked peptides that are consistently quantified in both label-swap replica are accepted. Optionally, cross-linked peptides that are quantified otherwise or not quantified can be re-quantified and validated using a semi-automated procedure, here Pinpoint.
4. The Pinpoint platform allows semi-manually correcting questionable results from MaxQuant. This process can be carried out for all or only a subset of cross-linked peptides of interests. Again, only cross-linked peptides that are consistently quantified in both label-swap replica are accepted.
5. Peptide results are combined to linked residue pairs, taking the median of supporting cross-linked peptides. A linked residue pair is unique to one conformation only if all its supporting data are unique to this conformation. Conflicts such as a cross-linked peptide that disagrees with other cross-linked peptides of the same linked residue pair can be revisited and validated manually. As this process requires high ethical conduct all raw data should be made available via a trusted public data repository such as ProteomeXchange.
6. Further structural interpretation is based on quantitation results of linked residue pairs.

**Figure 5:**
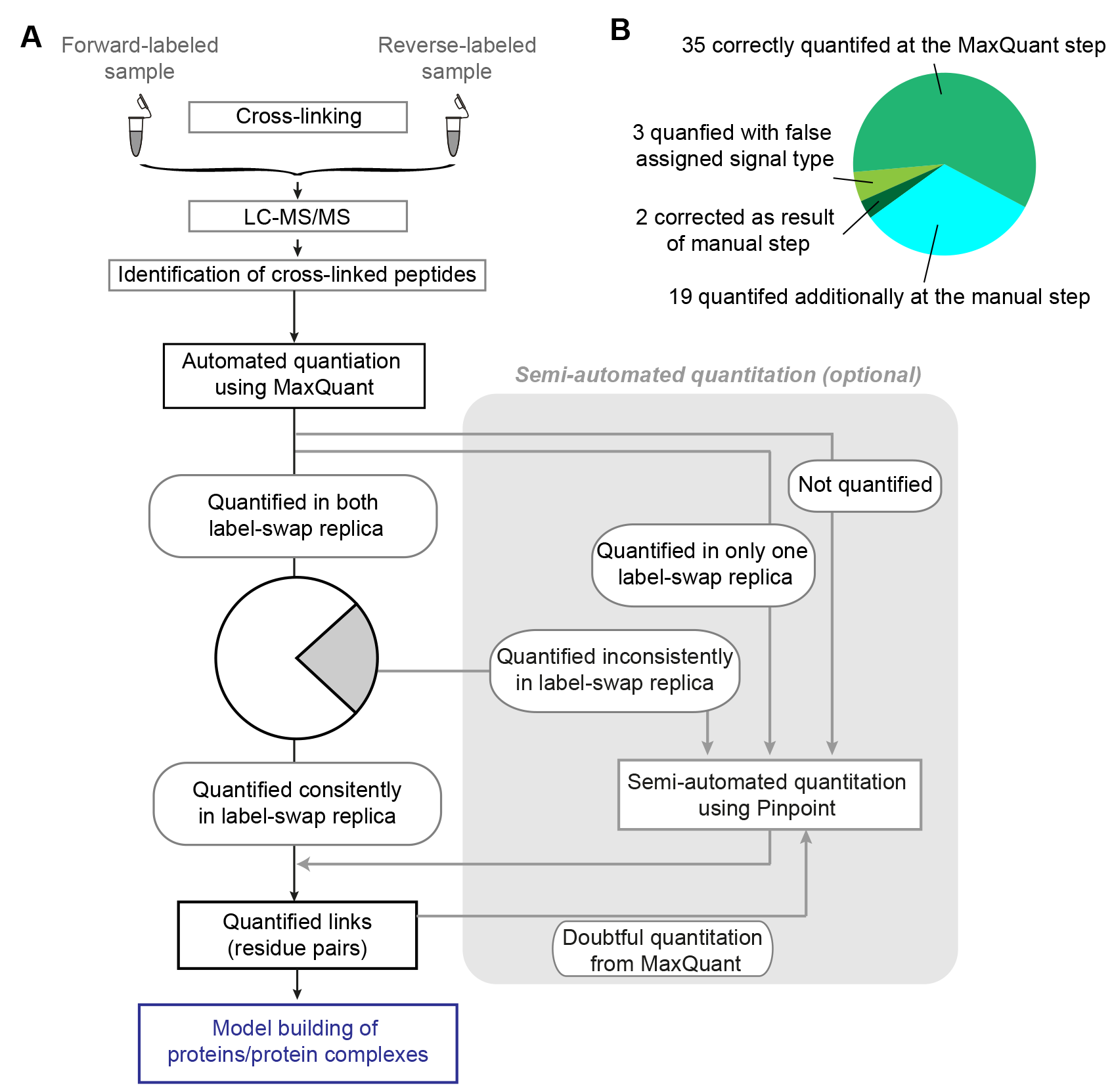
An integrate workflow for QCLMS analysis. (**A**) An integrated workflow for QCLMS including replicated experiments with label-swap and quantitation using Pinpoint and MaxQuant. (**B**) Summary of quantitation results on the reference dataset using the integrated workflow shown in (A).inn

We applied this data analysis process on our reference dataset (Fig. 5B). As we showed above, 66 cross-linked peptides were quantified by MaxQuant in forward and reverse replica. This identified and quantified 40 linked residue pairs and forms the final result of our automated quantitation. From the 59 residue pairs on our reference list MaxQuant achieved a recall rate of 68% (40 of 59) with an accuracy of 88% (35 of 40) in recognizing singlet and doublet peaks and an R^2^=0.93 for the quantification of doublets. This is the final result of an automated analysis by MaxQuant. In a second round we used Pinpoint only on those cross-linked peptides that were excluded from the results of the automated analysis because they were not consistently quantified in our label-swap replica. This resulted in an additional 39 confidently quantified cross-linked peptides. These added 19 new quantified residue pairs and led to correction for 2 of the 5 mis-classified doublet residue pairs of the automated process. Consequently, only 3 doublets remained singlets in the results of the extended workflow. In summary, we quantified all 59 expected linked residue pairs from our reference (100% recalls). 56 were classified correctly, yielding 96% accuracy.

The actual gain of the additional manual analysis will vary from experiment to experiment. In our reference dataset, dataset 2, automated quantitation using MaxQuant reliably quantified cross-links that are unique to either conformation: 14 of 16 C3 unique cross-links and 12 of 14 C3b unique cross-links were correctly quantified. These correctly quantified cross-links reflected all four major conformational changes in the alpha chain of C3 during the transition to C3b. This included 1) the exclusive existence of the ANA domain in C3 and its relative position in the molecule; 2) the relocation of the CUB-TED domain in C3b; 3) the reposition of the α'–N terminus in C3b and 4) the rearrangement of the shoulder region of the “shoulder” region of the molecule. Nine cross-links that were correctly quantified by MaxQuant as common in both C3 and C3b reflect the structural features that are similar between two conformations within individual domains and also between domains in the beta chain of the protein. Additional quantified/corrected cross-links using our semi-manual procedure are mainly (20 of 24) observed in both C3 and C3b. They provided a higher data density, confirming differences and similarities between structures of C3 and C3b revealed already by fully auto quantified dataset.

As an alternative to PinPoint (Thermo), software packages like Skyline (18) can be used. Also in Skyline, cross-linked peptides can be introduced by linearizing crosslinked peptide sequences in the same way as we established for Pinpoint (7). One caveat of the current version of Skyline is that it does not allow for grouping cross-linked peptide based on unique cross-linked residue pairs. As a consequence, the post-quantitation data processing becomes more elaborate.

## Conclusion

Our new release of MaxQuant (version 1.5.4.1) enables quantitation for cross-linked peptides. The underlying “Quantification only” mode of this new version of MaxQuant is not limited to the application for cross-linking data. Any analyte not identified within the MaxQuant workflow can now be quantified within MaxQuant. Detailed evaluation using a reference cross-link dataset showed that a fully automated process was subject to errors that are revealed by label-swopping. Requiring consistent quantitation in label-swap replicas significantly improved the quantitation accuracy. Manually assessing problematic peptide signals can improve both the recall rate and accuracy of dataset. Manual analysis is, however, time consuming and requires user expertise. The manual step is optional and might be applied selectively to data of interest. For example, interest might focus on a subset of cross-links between certain proteins in a large protein network, or a linkage pair that is key for drawing a structural conclusion. This integrated workflow shows best handling efficiency, recall rate, and quantitation accuracy. Opening MaxQuant to work with cross-links, introducing label-swap replica analysis to QCLMS and soliciting data sharing will hopefully help to consolidate quantitative cross-linking into a more routine approach.

## Acknowledgments

The Wellcome Trust generously funded this work through a Senior Research Fellowship to JR (103139), a Centre core grant (092076) and an instrument grant (108504). We also acknowledge the PRIDE team for the deposition of our data to the ProteomeXchange Consortium.

## References

1. Kalkhof, S., Ihling, C., Mechtler, K., and Sinz, A. (2005) Chemical cross-linking and high-performance Fourier transform ion cyclotron resonance mass spectrometry for protein interaction analysis: application to a calmodulin/target peptide complex. Anal Chem 77, 495–503

2. Mueller, D.R., Schindler, P., Towbin, H., Wirth, U., Voshol, H., Hoving, S., and Steinmetz, M. O. (2001) Isotope-tagged cross-linking reagents. A new tool in mass spectrometric protein interaction analysis. Anal Chem 73, 1927–1934

3. Pearson, K. M., Pannell, L. K., and Fales, H. M. (2002) Intramolecular crosslinking experiments on cytochrome c and ribonuclease A using an isotope multiplet method. Rapid Commun Mass Spectrom 16, 149–159

4. Rinner, O., Seebacher, J., Walzthoeni, T., Mueller, L. N., Beck, M., Schmidt, A., Mueller, M., and Aebersold, R. (2008) Identification of cross-linked peptides from large sequence databases. Nat Methods 5, 315–318

5. Fischer, L., Chen, Z. A., and Rappsilber, J. (2013) Quantitative crosslinking/mass spectrometry using isotope-labelled cross-linkers. Journal of proteomics 88, 120–128

6. Schmidt, C., Zhou, M., Marriott, H., Morgner, N., Politis, A., and Robinson, C. V. (2013) Comparative cross-linking and mass spectrometry of an intact F-type ATPase suggest a role for phosphorylation. Nature communications 4, 1985

7. Chen, Z. A., Fischer, L., Tahir, S., Bukowski-Wills, J.-C., Barlow, P. N., and Rappsilber, J. (2016) Quantitative cross-linking/mass spectrometry reveals subtle protein conformational changes. bioRxiv. dio:10.1101/055418

8. Tomko, R. J., Jr., Taylor, D. W., Chen, Z. A., Wang, H. W., Rappsilber, J., and Hochstrasser, M. (2015) A Single alpha Helix Drives Extensive Remodeling of the Proteasome Lid and Completion of Regulatory Particle Assembly. Cell 163, 432–444

9. Schmidt, C., and Robinson, C. V. (2014) A comparative cross-linking strategy to probe conformational changes in protein complexes. Nat Protoc 9, 2224–2236

10. Kukacka, Z., Rosulek, M., Strohalm, M., Kavan, D., and Novak, P. (2015) Mapping protein structural changes by quantitative cross-linking. Methods 89, 112–120

11. Cox, J., and Mann, M. (2008) MaxQuant enables high peptide identification rates, individualized p.p.b.-range mass accuracies and proteome-wide protein quantification. Nat Biotechnol 26, 1367–1372

12. Ong, S. E., Blagoev, B., Kratchmarova, I., Kristensen, D. B., Steen, H., Pandey, A., and Mann, M. (2002) Stable isotope labeling by amino acids in cell culture, SILAC, as a simple and accurate approach to expression proteomics. Mol Cell Proteomics 1, 376–386

13. Cox, J., Hein, M. Y., Luber, C. A., Paron, I., Nagaraj, N., and Mann, M. (2014) Accurate proteome-wide label-free quantification by delayed normalization and maximal peptide ratio extraction, termed MaxLFQ. Mol Cell Proteomics 13, 2513–2526

14. Beck, S., Michalski, A., Raether, O., Lubeck, M., Kaspar, S., Goedecke, N., Baessmann, C., Hornburg, D., Meier, F., Paron, I., Kulak, N. A., Cox, J., and Mann, M. (2015) The impact II, a very high resolution quadrupole time-of-flight instrument for deep shotgun proteomics. Mol Cell Proteomics 14, 2014–2029

15. Vizcaino, J. A., Deutsch, E. W., Wang, R., Csordas, A., Reisinger, F., Rios, D., Dianes, J. A., Sun, Z., Farrah, T., Bandeira, N., Binz, P. A., Xenarios, I., Eisenacher, M., Mayer, G., Gatto, L., Campos, A., Chalkley, R. J., Kraus, H. J., Albar, J. P., Martinez-Bartolome, S., Apweiler, R., Omenn, G. S., Martens, L., Jones, A. R., and Hermjakob, H. (2014) ProteomeXchange provides globally coordinated proteomics data submission and dissemination. Nat Biotechnol 32, 223–226

16. Cox, J., Neuhauser, N., Michalski, A., Scheltema, R. A., Olsen, J. V., and Mann, M. (2011) Andromeda: a peptide search engine integrated into the MaxQuant environment. J Proteome Res 10, 1794–1805

17. Kustatscher, G., Hegarat, N., Wills, K. L., Furlan, C., Bukowski-Wills, J. C., Hochegger, H., and Rappsilber, J. (2014) Proteomics of a fuzzy organelle: interphase chromatin. EMBO J 33, 648–664

18. MacLean, B., Tomazela, D. M., Shulman, N., Chambers, M., Finney, G. L., Frewen, B., Kern, R., Tabb, D. L., Liebler, D. C., and MacCoss, M. J. (2010) Skyline: an open source document editor for creating and analyzing targeted proteomics experiments. Bioinformatics 26, 966–968

